# Distinct Brain Systems Support Afferent and Efferent Autonomic Activity

**DOI:** 10.64898/2026.06.03.729710

**Authors:** Jungwon Min, Gengshuo Liu, Martin J. Dahl, Tae-ho Lee, Kaoru Nashiro, Hyun Joo Yoo, Christine Cho, Paul Bogdan, Catie Chang, Paul Lehrer, Julian F. Thayer, Mara Mather

**Affiliations:** University of Southern California, Los Angeles, CA, US; Max Planck Institute for Human Development, Berlin, BE, DE; Virginia Polytechnic Institute and State University, Blacksburg, VA, US; Vanderbilt University, Nashville, TN, US; Rutgers Robert Wood Johnson Medical School, New Brunswick, NJ, US; University of California Irvine, Irvine, CA, US

**Keywords:** emotion regulation, resting state, heart rate variability, afferent, efferent, central autonomic network (CAN)

## Abstract

Numerous studies have reported brain correlates of autonomic activity. However, ambiguity exists regarding whether those correlates reflect receiving or sending out autonomic signals. Additionally, little is known about how emotion and aging interact with brain-autonomic coordination. Based on time-varying heart rate variability (HRV) and blood oxygenation level dependent (BOLD) data (N = 104 younger; N = 51 older adults), we found insular and cingulate activity was negatively correlated with HRV during emotion regulation. We further examined the afferent and efferent nature of HRV-BOLD correlations during rest (N = 102 younger; N = 51 older adults). Functionally afferent regions where decreases in HRV were associated with subsequently increased BOLD activity included the posterior insula, postcentral gyrus and frontal pole. Functionally efferent regions where increased activity was associated with subsequently increased HRV included the anterior insula and cingulate. Information appeared to flow in one direction as the afferent regions’ activity Granger-predicted the efferent regions’ activity. Together, our findings suggest a feedback loop where decreased HRV increases the afferent regions’ activity, which activates the efferent regions, leading to increased HRV. Aging seems to affect this system; the efferent and afferent regions scarcely overlapped for younger adults, but did overlap for older adults.

## Introduction

Emotion and autonomic activity are coupled. The central autonomic network (CAN) coordinates emotional behavior and autonomic signals (Craig, 2003; Hagemann et al., 2003; Satpute et al., 2019; Siegel et al., 2018). While anatomical studies have located nodes of the CAN in the human brain (Benarroch, 2020), how the identified regions interact with each other to represent or control emotional behavior is as yet unclear (Barrett and Simmons, 2015; Craig, 2002; Critchley and Harrison, 2013; Damasio and Carvalho, 2013; Kleckner et al., 2017). One uncertainty lies in whether previously identified regions are receiving incoming information from or sending out commands to the periphery (Candia-Rivera et al., 2022; Chouchou et al., 2019; Critchley, 2005; Critchley et al., 2004; Khalsa et al., 2009; Oppenheimer and Cechetto, 2016). Direct stimulation studies with implanted electrodes in the human brain may provide more straightforward findings. However, even with direct stimulation, inconsistencies still exist. For example, the insula is known as the main substrate for regulating autonomic signals (Craig, 2002), but some studies do not agree on its subregional roles (Chouchou et al., 2019; Oppenheimer and Cechetto, 2016) and a lesion study suggested that the insula might not be critical for heartbeat sensation (Khalsa et al., 2009). In addition, those studies are based on a small number of neurosurgical patients with a priori sets of regions such as the insula, anterior cingulate cortex (ACC), and amygdala (Gillies et al., 2019; Inman et al., 2020; Oppenheimer et al., 1992) and provide limited interpretations about their interactions.

Functional magnetic resonance imaging (fMRI) studies have developed a variety of methods to capture the cortical and subcortical representation of autonomic regulation (Beissner et al., 2013; Critchley, 2005; Ruiz Vargas et al., 2016; Thayer et al., 2012). Studies employed cognitive, emotional, or physical tasks that could cause changes in peripheral autonomic activity, and related the evoked changes to brain blood oxygenation level dependent (BOLD) activity (Critchley, 2005). To locate autonomic control regions, researchers developed tasks involving physical and mental arousal and attributed the task-induced shifts in physiological states (e.g., heart rate or blood pressure) to the changes in brain activity (Coulson et al., 2015; Critchley et al., 2005b, 2003, 2000; Fechir et al., 2010; Gianaros et al., 2004; Matsunaga et al., 2009; Napadow et al., 2008). The brain representation of incoming autonomic signals was also examined by using task scans, during which participants attended to internal sensations such as their own heartbeats and arousal states (Critchley et al., 2004; Goswami et al., 2011; Pollatos et al., 2007) or experienced perturbations in the baroreflex which controls heart rate to maintain steady blood pressure (Basile et al., 2012; Kimmerly et al., 2005). Whereas multiple tasks were designed to locate autonomic brain regions, tasks investigating autonomic regions associated with emotional processes are scarce (Critchley et al., 2005a, 2000; Garfinkel et al., 2015; Gray et al., 2012; Harrison et al., 2010; Lane et al., 2009).

The fMRI studies investigating brain areas associated with afferent or efferent autonomic signals converge on a few common regions (e.g., insula and ACC; Beissner et al., 2013; Critchley, 2005; Oppenheimer and Cechetto, 2016; Ruiz Vargas et al., 2016). This raises questions as to 1) whether afferent and efferent activities are controlled by those shared brain regions and 2) whether there are alternative methods to disentangle the two activities. Challenges come from anatomical arrangements where afferent and efferent activities interact at multiple levels involving reflexes, brainstem modulation, and higher cortical functioning (Benarroch, 2016, 1993). In fact, tasks designed to increase afferent autonomic activity often demand mental effort, which can also activate multiple effort-induced arousal (efferent) pathways and obscure the identification of the afferent regions (Critchley, 2005; Ziegler et al., 2009).

Resting-state data without effortful behavior might help bypass the limitations of the task-based fMRI studies. Some investigators set cardiovascular activity as a regressor of interest and identified covarying brain regions during resting-state fMRI (Ziegler et al., 2009; Chang et al., 2009, 2013; Golestani et al., 2016; Tong and Frederick, 2014; Valenza et al., 2020). These studies suggest that the default mode network, ACC, amygdala, ventromedial prefrontal cortex (PFC) and posterior cingulate cortex (PCC) are associated with cardiovascular activity. While the studies developed new analysis techniques and located the CAN regions, they did not address whether those brain regions receive afferent signals or generate efferent signals. To disentangle afferent and efferent activities in the brain, we propose a regression model which associates brain BOLD activity with physiological activity within two types of separate 10-second time windows positioned before and after each brain volume (Figure 1). For example, by correlating BOLD signals with physiological activity within time windows positioned before brain volumes (while accounting for physiological activity within time windows positioned after brain volumes), we can locate brain regions associated with afferent activity.

**Figure 1.**
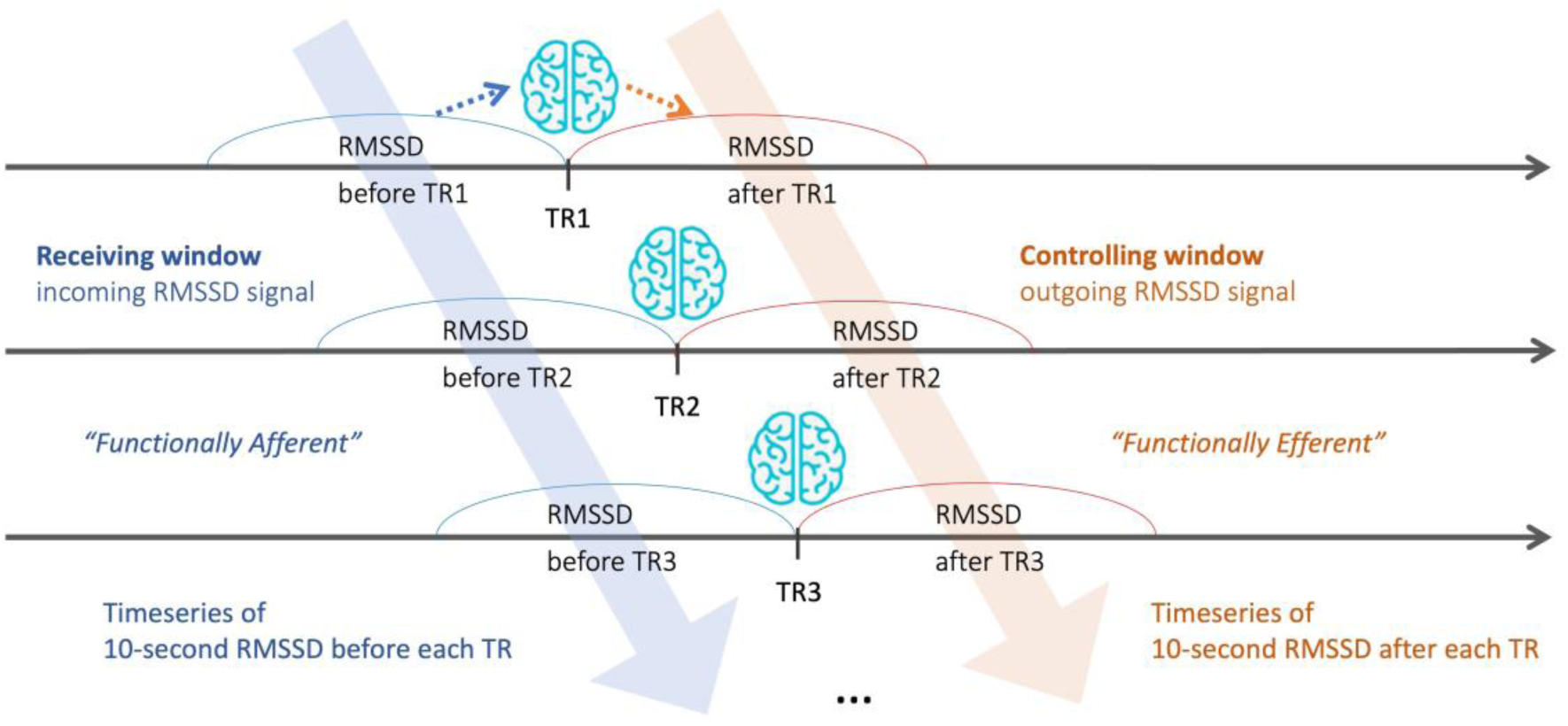
Two types of RMSSD time series paired with whole-brain BOLD signals. Modeling individual BOLD time series at TR with RMSSD time series over two 10-second windows before and after TRs allows us to investigate bidirectional relationships between RMSSD and brain BOLD activity. By correlating at-TR BOLD signals with before-TR RMSSD while accounting for after-TR RMSSD, we can examine how changes in RMSSD influence changes in BOLD activity. Likewise, by correlating at-TR BOLD signals with after-TR RMSSD while accounting for before-TR RMSSD, we can investigate how changes in RMSSD are influenced by changes in BOLD activity.

Aging adds to the complexity of disentangling afferent and efferent signaling. With aging, sympathetic activity increases and parasympathetic activity decreases (Mather, 2024; Seals and Esler, 2000; Zulfiqar et al., 2010). Diminished integrity of the afferent network or waning effectiveness of the efferent network may cause these autonomic shifts. Aging brains typically have reduced BOLD activity, show less specific response to task conditions, and show less segregation between networks at rest (Cabeza, 2001; Deery et al., 2023; Hu et al., 2014; Huettel et al., 2001). Examining brain BOLD activity in both younger and older adults may provide clues regarding age-related changes in afferent and efferent CAN activity.

While prior literature suggests heart rate variability (HRV) measured during rest as a biomarker of emotion regulation ability (Thayer and Lane, 2009), findings on HRV measured during tasks are not consistent and did not consider the time-varying nature of HRV (Butler et al., 2006; Lane et al., 2009; Wawrzyniak et al., 2016). Thus, we first investigated the brain correlates of time-varying HRV during emotion regulation by using generalized psychophysiological interaction analysis (McLaren et al., 2012). We then examined whether the brain regions associated with time-varying HRV during emotion regulation are involved with either afferent or efferent activity by using resting-state data. As afferent regions are related to incoming autonomic signals, we hypothesized afferent regions would overlap with previously reported interoceptive regions within the insula, ACC, and somatosensory areas (Adolfi et al., 2017; Critchley et al., 2004; Pollatos et al., 2007). As efferent regions are associated with outgoing autonomic signals, we hypothesized that efferent regions would overlap with motor command pathways involving the ACC, anterior insula, thalamus, and amygdala (Critchley et al., 2005b; Williamson et al., 2002). Thus, we predicted distinct brain regions for afferent and efferent autonomic activities. In addition, we predicted that this distinctiveness will be reduced for older adults as brain activity becomes less segregated with aging (Cabeza, 2001; Deery et al., 2023).

## Methods

### Participants

We used functional magnetic resonance imaging (fMRI) data collected at the pre-intervention phase of a HRV biofeedback intervention study (ClinicalTrials.gov Identifier: NCT03458910), approved by University of Southern California’s Institutional Review Board (Nashiro et al., 2022; Yoo et al., 2023). Younger and older participants without serious medical or psychiatric illness were recruited via USC’s subject pool, USC’s online bulletin board, Facebook, and flyers. Older adults with TELE scores lower than 16 were excluded from enrollment (Gatz et al., 1995). We obtained informed consent from all the participants and paid them $15 per hour for their lab visits. We used data from 104 younger adults and 51 older adults for the emotion regulation task (Table 1). We used resting-state scan data from 153 participants, 102 younger adults and 51 older adults (Table 1).

**Table 1.** Age, education, and sex of participants. *Note.* The original intervention study recruited 193 adults, among whom 172 participants provided emotion-regulation task scans and 176 participants provided resting-state scans at their pre-intervention phase. From the emotion regulation task data, we excluded 5 participants whose multi-echo independent component analysis (me-ICA) preprocessing pipeline failed and 12 participants whose photoplethysmography (PPG) data were not available or highly noisy. From the resting-state data, we excluded two participants whose me-ICA preprocessing pipeline failed and 21 participants whose PPG data were not available or contained high levels of noise. The number of older adults is lower than that of younger adults because data collection was halted due to COVID-19.

| Dataset | Age group | N (N males) | Mean Age (SD) | Min - Max | Mean Education (SD) | Min - Max |
| --- | --- | --- | --- | --- | --- | --- |
| Emotion Regulation | Younger | 104 (53 males) | 22.8 (2.76) | 18 - 31 | 16.0 (2.12) | 12 - 24 |
|  | Older | 51 (19 males) | 65.06 (6.69) | 55 - 80 | 16.7 (2.31) | 12 - 25 |
| Resting State | Younger | 102 (53 males) | 22.78 (2.77) | 18 - 31 | 16.0 (2.15) | 12 - 24 |
|  | Older | 51 (19 males) | 65.02 (6.35) | 55 - 80 | 16.87 (2.36) | 13 - 25 |

### MRI data acquisition

We collected MRI data at USC’s Dana and David Dornsife Cognitive Neuroimaging Center using a 3T Siemens MAGNETOM Prisma MRI scanner with a 32-channel head coil (Yoo et al, 2023). We collected 175 whole-brain volumes for resting-state scan and 250 for emotion-regulation task scan of T2*-weighted functional images using a multi-echo planar imaging sequence [repetition time (TR) = 2,400 ms, echo time (TE) = 18/35/53 ms, slice thickness = 3.0 mm, flip angle = 75°, field of view = 240 mm, voxel size = 3.0 mm isotropic]. We then obtained a T1-weighted MPRAGE anatomical image [TR = 2,300 ms, TE = 2.26 ms, slice thickness = 1.0 mm, flip angle = 9°, field of view = 256 mm, voxel size = 1.0 mm isotropic] to aid coregistration.

### Emotion regulation task

The task was based on a previously published emotion regulation task where participants used cognitive reappraisal strategies to up- or down-regulate emotion for positive and negative emotions (Kim and Hamann, 2007). The main findings of the task are published (Min et al., 2024, 2022). The task lasted for 10 minutes with 12 blocks. Each block contained three trials (Figure S1). Each trial consisted of three phases: instruction, regulation, and rating (Min et al., 2022). During the 1-second instruction, participants were presented with the words “intensify,” “diminish,” or “view” on a black screen. During the following 6-second regulation phase, they regulated or passively experienced emotions elicited by the presented images, which were negative, positive, or neutral. During the final 4-second rating phase, they rated the strength of their feeling in the moment (levels of emotional intensity) by pressing a button of 1 to 4.

### HRV data acquisition and processing

We used PPG signals rather than electrocardiogram (ECG). We found that PPG is less susceptible to radio frequency interference than ECG during MRI and previous work indicates it shows strong correlations in heart beat measures with ECG (Kumar et al., 2021; Lu et al., 2009; Schumann et al., 2021; Weinschenk et al., 2016). Heartbeat data was collected at 10 kHz sampling rate concurrently with the functional MRI scans using the PPG sensor attached on the left middle finger. The PPG data was down-sampled to 1 KHz and entered into Kubios HRV Premium 3.1 software with automatic beat correction to compute beat-to-beat intervals for each participant. The extracted intervals were used to calculate the root mean square of successive differences (RMSSD) between the intervals over 10-second time windows using Matlab (R2021b). We chose to calculate RMSSD over a series of sliding temporal windows to capture the time-varying nature of parasympathetic activity (Krause et al., 2023; Shaffer and Ginsberg, 2017) as, compared with frequency-domain measures, RMSSD is less influenced by respiration and can be computed with shorter periods of data (Shaffer and Ginsberg, 2017).

During the emotion regulation task, the RMSSD series was computed over 10-second time windows centered around each TR and entered as a physiological regressors to a psychophysiological interaction analysis. For resting-state scans, the afferent type of RMSSD series (before-TR RMSSD) was computed over time windows that began 10 seconds before each TR and ended at each TR (Figure 1). The efferent type of RMSSD series (after-TR RMSSD) was computed over time windows that began at each TR and ended 10 seconds after each TR. For the within-subject means and ranges of before-TR and after-TR RMSSD series see Table S1 and for their correlations see Figure S2.

### MRI data analysis

For both the emotion-regulation and resting-state data, we used the me-ICA preprocessing pipeline to form a single time series across three echos and to remove artifact components based on the linear echo-time dependence of BOLD signal fluctuations (Kundu et al., 2013, 2012). After denoising the fMRI data, we used FMRIB Software Library (FSL) version 6.0 for the individual- and group-level analyses (Jenkinson et al., 2012; Woolrich et al., 2004, 2001). We accounted for the hemodynamic delay of BOLD signal relative to PPG signal by shifting the BOLD data 5-second backward compared to the PPG data (Chang et al., 2013). During the individual-level analysis, each functional image was spatially smoothed with a Gaussian kernel of full width at half maximum 5mm, motion corrected with MCFLIRT, registered to the MNI152 T1 2mm template via its T1-weighted anatomical image using a 12-degree of freedom affine linear transformation with FLIRT, and prewhitened with FILM. High-pass filtering with 600-second temporal width was applied to the emotion regulation data.

To find brain regions which correlate with RMSSD during emotion regulation, we ran generalized psychophysiological interaction analysis (McLaren et al., 2012) using FSL, where individual whole-brain BOLD signals were modeled with fifteen regressors: the seven psychological regressors of no interest, one physiological regressor (RMSSD), and seven psychophysiological interaction terms between the seven psychological regressors and the physiological regressor. The seven psychological regressors were the 36-second block periods of task conditions (diminish-negative, view-negative, intensify-negative, diminish-positive, view-positive, intensify-positive, and view-neutral), each of which was convolved with a double-gamma hemodynamic response function. The physiological regressor is the series of RMSSD averaged over 10-second periods centered around each TR. The interaction term was the multiplication of the psychological and physiological regressors. The final results were based on FSL’s mixed-effects model (FLAME 1) including younger and older adults and were corrected for family-wise error at *p* < .05 with the cluster-size threshold at *Z* > 3.1.

We analyzed the resting-state data to examine the afferent and efferent nature of the clusters correlated with RMSSD during the emotion regulation task. We paired the whole-brain volume at each measurement (TR), with two non-overlapping 10-second time windows: the one ending at the TR and the other starting at the TR (Figure 1). We assumed that RMSSD values obtained from time windows starting 10 seconds before and ending at every TR (before-TR RMSSD) contained more incoming cardiac signals to the brain. Likewise, we assumed that RMSSD values from time windows starting at every TR and ending 10 seconds after (after-TR RMSSD) carried more outgoing brain signals to the heart. By modeling brain BOLD time series with before-TR and after-TR RMSSD time series, we examine ‘functionally afferent’ and ‘functionally efferent’ CAN regions respectively. For brevity of notation, we label regions correlated with before-TR RMSSD as ‘afferent’ and labeled regions correlated with after-TR RMSSD as ‘efferent’ in the current study. To find the brain correlates of one type of RMSSD time series while accounting for the other type, we included the before-TR and after-TR RMSSD time series in one model. The two types of regression parameter estimates were averaged for younger and older adults by using FSL’s mixed-effect model (FLAME 1) with cluster-wise correction (*Z* > 3.1, *p* < 0.05). We followed up with some sensitivity analyses to confirm the results are HRV-specific (Figure S3) and are not biased by regressor collinearity (Figure S4), support the negative feedback loop idea (Figure S5), and are robust with covariates (Figure S6). As we noticed the thalamus occupying large parts of both the afferent and efferent clusters (Figure 3A), we examined the nature of the activated thalamic regions (Figure S7).

As post-hoc analyses of the relationship between the afferent and efferent brain regions in the resting-state data, we calculated cross-correlations between the two afferent and efferent regions’ BOLD time series for each participant. We first binarized the afferent and efferent regions (blue and orange in Figure 3) and excluded their overlapping regions (red in Figure 3). Using the two masks separately for younger and older adults, we extracted BOLD time series, normalized them within each participant, and calculated their cross-correlations with time lags from -4TR to +4TR. Four TRs (9.6 seconds) approximate the length of the time window (10 seconds) within which we calculated RMSSD. We first ran one-sample *t*-tests, contrasting the cross-correlation coefficient against zero for each time point, separately for younger and older adults (9 comparisons for each age group). For this, we Bonferroni-corrected *p* values for a total of 18 comparisons.

As we noticed an asymmetry in the cross-correlation plot for younger adults only (Figure 5A), we compared the mean skewness of individual cross-correlation plots between younger and older adults. The skewness was estimated by fitting a skew normal distribution (𝛼, 𝜇, 𝜎; Mudholkar & Hutson, 2000) to the cross correlation values of each participant using the Curve Fitting Toolbox of Matlab (2023b). To better approximate the positive distribution function to the cross-correlation values ranging -1 to 1, we modified the function by adding a constant term (𝜔) to the original skew normal distribution. Assuming more associations for temporally close signals, we used the weighted nonlinear least squares method such that the afferent and efferent signals with less time lags were more weighted. After fitting the modified skew normal distribution curve to each participant’s cross correlation values over 9 TR points, we extracted the skewness parameter (𝛼) for each participant and ran a *t*-test to compare the means of 𝛼 between younger and older adults.

We further tested whether BOLD activity in the afferent regions predicts activity in the efferent regions or the other direction by employing the Granger causality test (Granger, 1969; Sargent et al., 2024). For this, we first ran the Granger causality analysis on the BOLD time series used for the cross-correlation analysis with both afferent-to-efferent and efferent-to-afferent directions. We log-transformed and ran a *t*-test on the Granger *F* statistics to compare the afferent-to-efferent predictability to the efferent-to-afferent predictability for younger and older adults.

## Results

To investigate the brain regions whose BOLD activity interacts with time-varying RMSSD during emotion regulation, we used generalized psychophysiological interaction analysis. The results revealed that throughout the whole task scan, RMSSD was negatively correlated with BOLD activity in the ACC, anterior insula, posterior insula, lingual gyrus, thalamus, putamen, and amygdala for younger adults (Figure 2). These areas increased their activity when RMSSD was decreased. Other regions also showed negative correlations with RMSSD during emotion up- and down-regulation (Figure S8).

**Figure 2.**
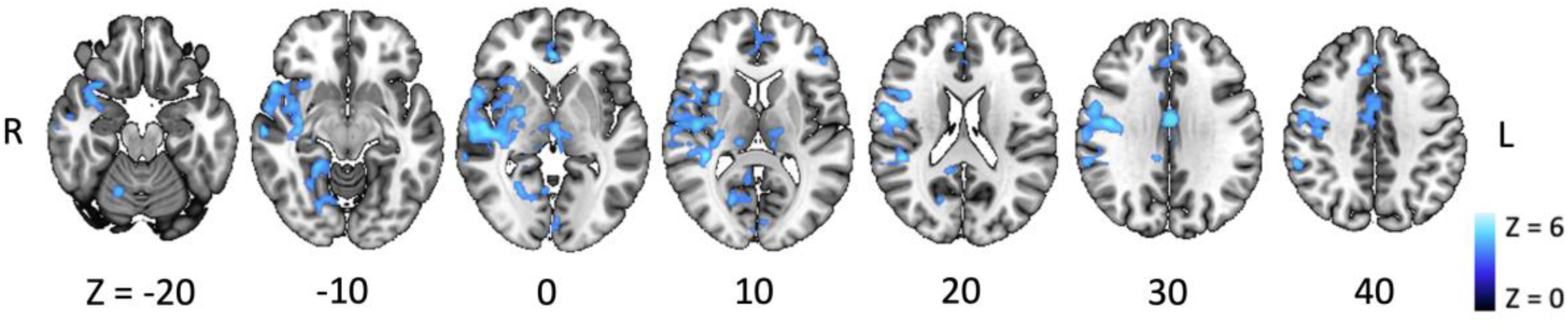
Brain regions associated with time-varying RMSSD during emotion regulation. Throughout the whole scan, BOLD activity in the ACC, anterior insula, and posterior insula showed negative correlations with RMSSD for younger adults. Neither negative nor positive correlations were found for older adults.

To locate functionally afferent brain regions, we correlated BOLD time series with before-TR RMSSD time series while accounting for after-TR RMSSD. For younger adults, we found negative correlations between before-TR RMSSD and BOLD activity in regions including the frontal pole, posterior insula, postcentral gyrus, central opercular cortex, superior temporal gyrus, paracingulate gyrus, thalamus, caudate, putamen, ACC, and PCC (Figure 3A, blue). Negative correlations for older adults were focused on smaller-sized clusters surrounding the frontal operculum cortex, posterior insula, precentral gyrus, postcentral gyrus (Figure 3B, blue). Neither younger nor older adults showed significant clusters with positive correlations between before-TR RMSSDs and BOLD activity. Compared with younger adults, older adults showed greater negative correlations between before-TR RMSSD and BOLD activity in the postcentral gyrus and frontal operculum cortex (Figure 4B, blue).

**Figure 3.**
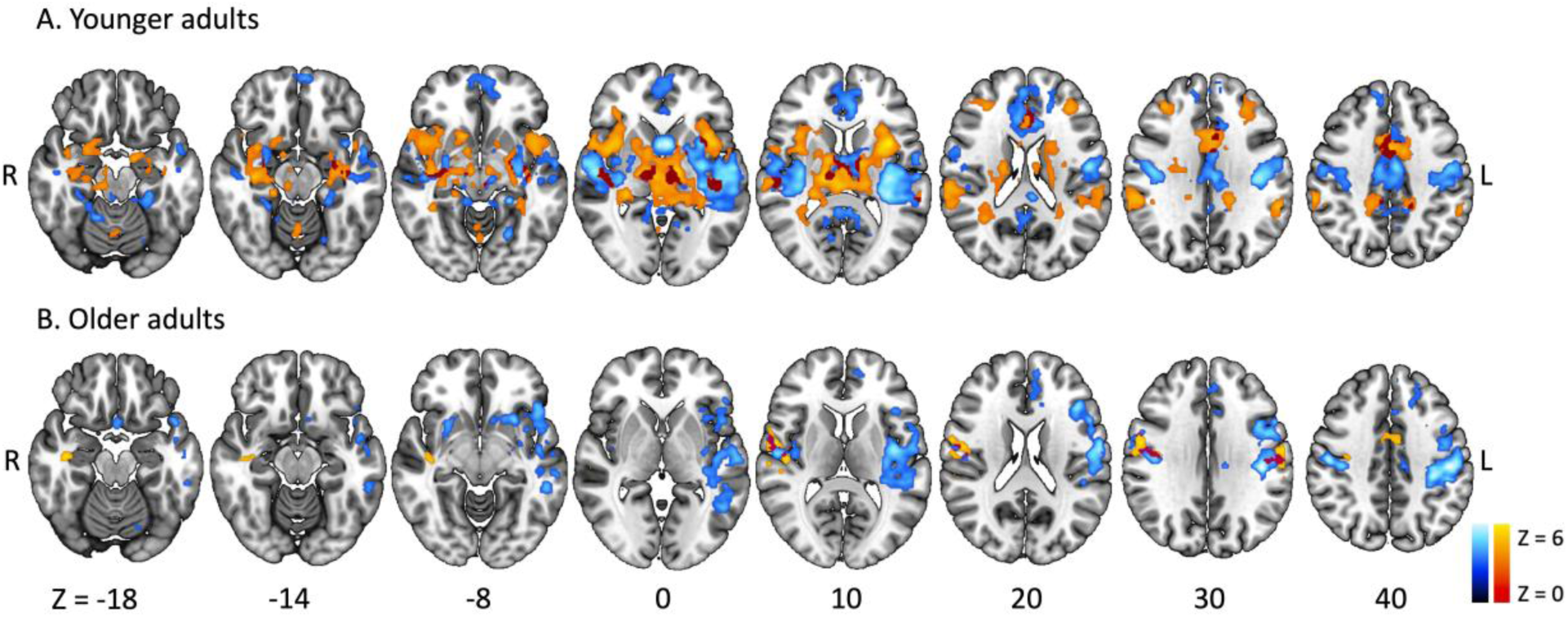
Functionally afferent and efferent brain regions for younger (A) and older adults (B). Blue clusters were negatively correlated with before-TR RMSSDs, and orange clusters were positively correlated with after-TR RMSSDs. The dark red color indicates the overlap between blue and orange clusters.

**Figure 4.**
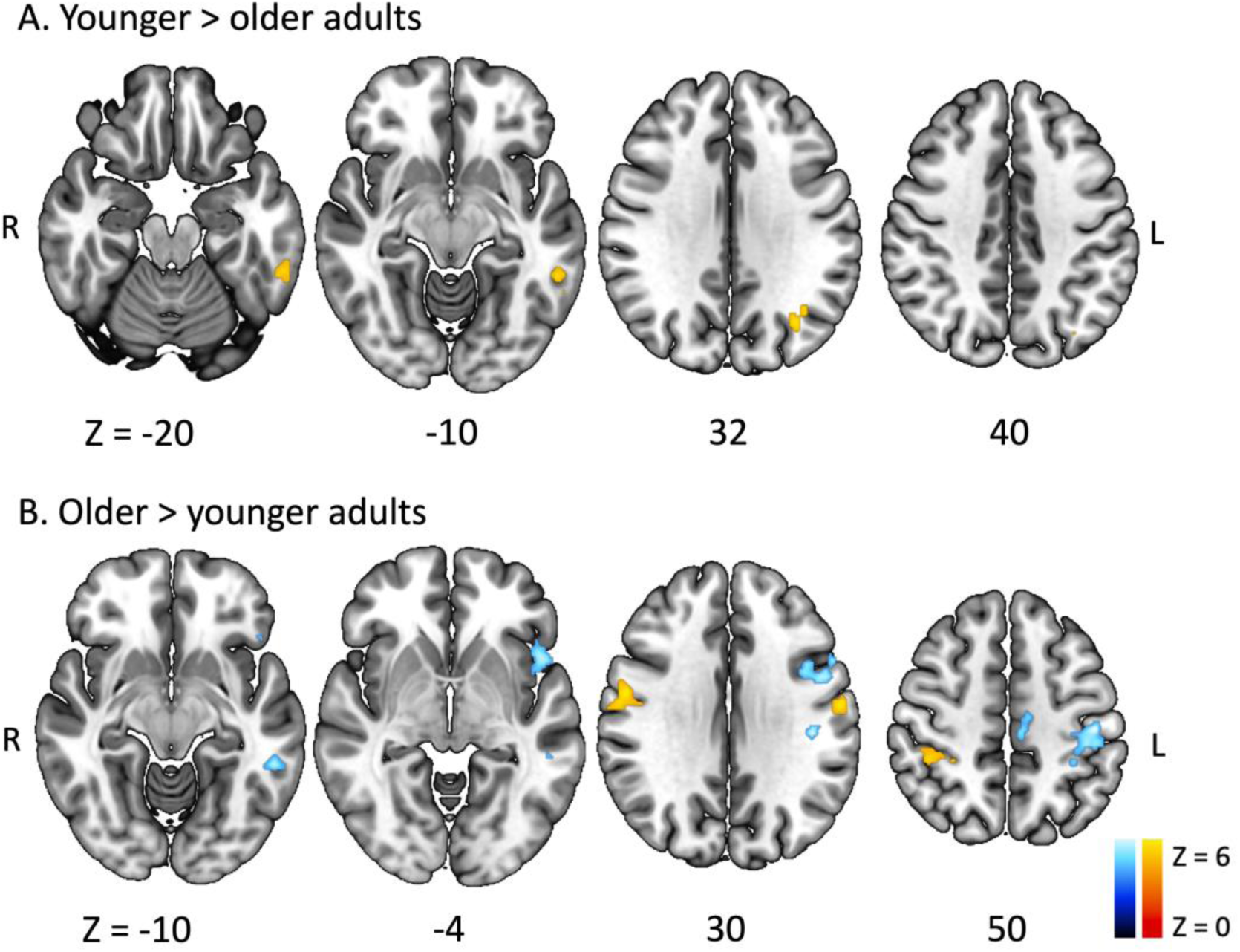
Difference between younger and older adults in afferent and efferent regions. Blue clusters were negatively correlated with before-TR RMSSDs, and orange clusters were positively correlated with after-TR RMSSDs.

To locate functionally efferent brain regions, we correlated BOLD time series with after-TR RMSSD time series while accounting for before-TR RMSSD. For younger adults, after-TR RMSSD was positively correlated with BOLD activity in the dorsal ACC, paracingulate gyrus, anterior insula, central opercular cortex, thalamus, putamen, hippocampus, brainstem, and cerebellum vermis (Figure 3A, orange). For older adults, a positive correlation with after-TR RMSSD was focused on the dorsal ACC, supplementary motor cortex, postcentral gyrus, and central opercular cortex (Figure 3B, orange). Neither younger nor older adults showed significant clusters with negative correlations between after-TR RMSSD and BOLD activity. Compared with older adults, younger adults showed a greater correlation between the after-TR RMSSD and BOLD activity in the left lateral occipital cortex, left inferior temporal gyrus, and cerebellum (left crus II) (Figure 4A, orange). Compared with younger adults, older adults showed greater correlation between after-TR RMSSD and BOLD activity in the precentral and postcentral gyrus (Figure 4B, orange). The afferent regions for younger adults covered broader spatial areas than the efferent regions, and the two types of regions did not overlap except a small portion, focused on the thalamus (Figure 3A, red). While the number of voxels for the afferent regions was 20,797, the number of voxels for the efferent regions was 15,406. The common area between the two regions had 1,368 voxels, 6.6% of the afferent region and 8.9% of the efferent region. For older adults, the afferent, efferent, and overlapping regions (Figure 3B, red) were diminished in volume (14,340, 1,743, and 371 voxels respectively). The overlapping regions were 2.6% of the afferent region and 21.3% of the efferent regions.

To further examine how the afferent regions influence efferent regions (Figure 3), we first ran cross-correlation analyses. We found positive cross-correlations between the two BOLD time series from the afferent and efferent regions. For younger adults (Figure 5A), all the cross-correlation coefficients were significant after correcting for the multiple comparisons (*t*s > 7; corrected *p*s < 0.001). For older adults (Figure 5A), the cross-correlation coefficients within 2 TR (4.8 seconds) timelags were significant (*t*s > 4; corrected *p*s < 0.001) along with one coefficient at 3 TR timelags (*t* = 3.29, corrected *p* = 0.04). By fitting the skew normal distribution with a constant term to the cross-correlation values, four parameters (𝛼, 𝜇, 𝜎, 𝜔) were estimated and their goodness of fit was measured by *r*^2^. Their means (standard errors) were respectively -1.43 (0.49), -0.08 (0.14), 26.21 (23.51), 0.23 (0.03) with *r*^2^ = 0.78 for younger adults and 0.23 (0.66), 0.02 (0.18), 2.41 (0.33), 0.20 (0.05) with *r*^2^ = 0.80 for older adults. We found a significant difference in skewness, 𝛼, between younger and older adults, *t*(151) = -2.0, *p* = 0.02. While younger adults showed strong negative skewness (M = -1.43, SE = 0.49), older adults showed positive skewness (M = 0.23, SE = 0.66).

**Figure 5.**
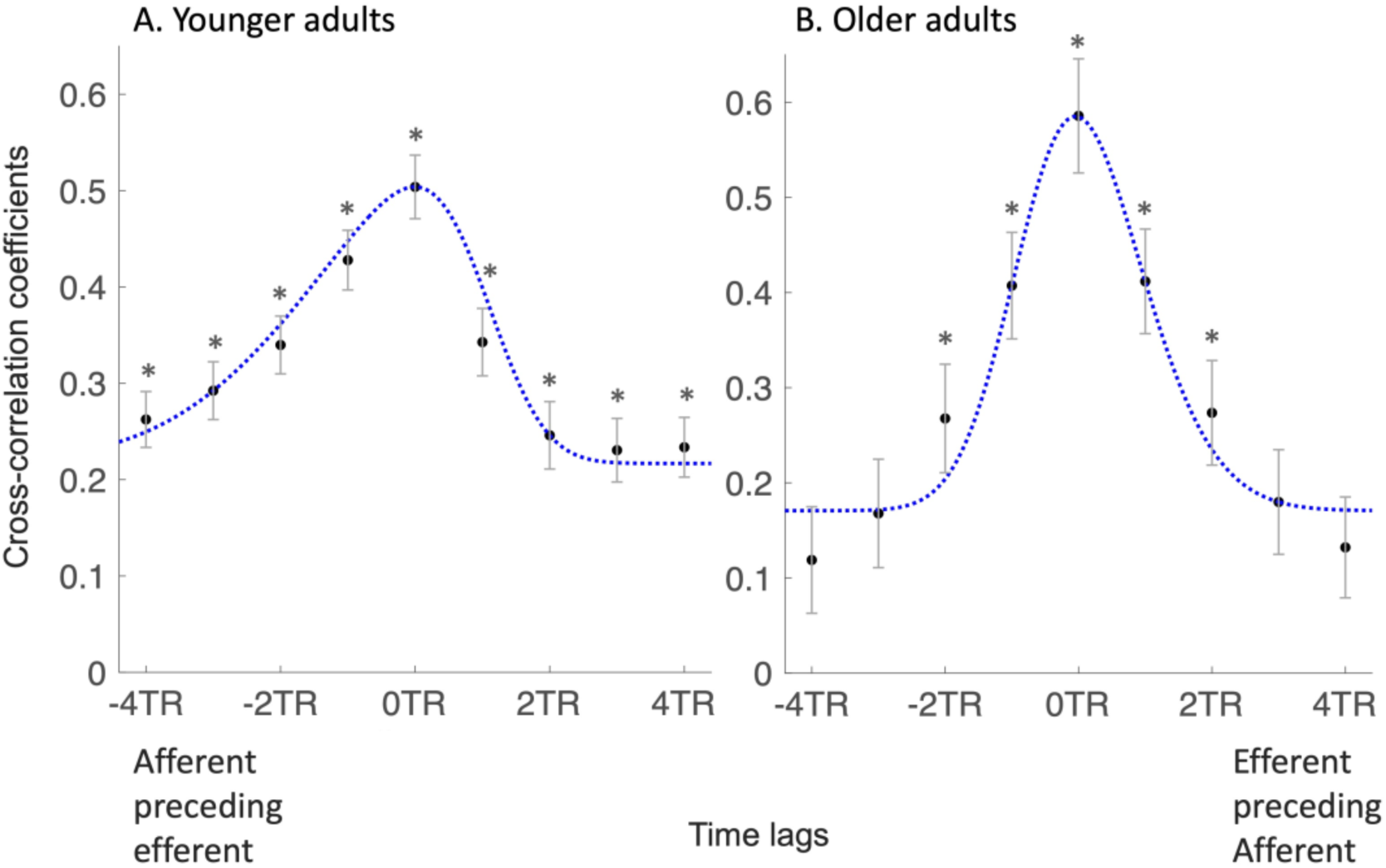
Cross-correlations between BOLD series from afferent and efferent regions. On the X axis, negative values (e.g., -4TR or -9.6 seconds) mean that in the analysis the afferent BOLD series precede the efferent BOLD series by the value (e.g., by 4 TRs or 9.6 seconds), and positive values (e.g., +2TR or +4.8 seconds) mean the efferent BOLD series precede the afferent BOLD series (e.g., by 2 TRs or 4.8 seconds). Cross-correlation coefficients at different time lags were plotted with standard error bars, and their significance (corrected *p* < 0.001) was indicated by asterisks. The dotted blue line is the skew normal distribution curve fitted to the mean cross-correlation coefficients separately for younger and older adults.

With the same BOLD time series from the afferent and efferent regions (Figure 3), we tested whether one type could predict the other. For younger adults, 59% of the participants showed significant Granger causality from the afferent-to-efferent direction, but 39% showed significant Granger causality for the opposite efferent-to-afferent direction. The *F*-statistics of Granger causality were greater for the afferent-to-efferent direction than the opposite direction for younger adults, *t*(101) = 3.94, *p* = 0.0001. For older adults, 59% showed significant Granger causality for the afferent-to-efferent direction, and similarly 57% showed significant Granger causality for the efferent-to-afferent direction. The *F*-statistics of Granger causality did not differ between the two directions for older adults, *t*(50) = -1.76, *p* = 0.09.

## Discussion

The current study investigated the brain correlates of time-varying RMSSD during emotion regulation and used resting-state data to examine which brain regions are associated with afferent and efferent RMSSD signals. As RMSSD measures successive differences in beat-to-beat intervals, it reflects relatively rapid changes in heart rate. Sympathetic nerve activity takes longer than parasympathetic vagus nerve activity to change heart rate (Elghozi and Julien, 2007) and so RMSSD is modulated by parasympathetic activity but shows little influence of sympathetic activity (Polanczyk et al., 1998). Throughout the emotion regulation task, the anterior insula, posterior insula, and ACC increased BOLD activity when RMSSD decreased (Figure 2). While the posterior and anterior insula are involved with receiving visceral signals and representing interoceptive information respectively, the cingulate cortex is more associated with modulating emotional and cardiovascular arousal (Craig, 2009; Critchley et al., 2004, 2003). At a glance, the brain correlates of RMSSD encompass two seemingly opposing roles: processing incoming signals and controlling outgoing signals. Since the emotion regulation task requires both interoceptive and arousal-modulating abilities (Min et al., 2022), BOLD activity during the task are likely to carry both afferent and efferent signals. By modeling resting-state BOLD signals with RMSSD values within 10-seconds before and after the time of every BOLD brain volume acquisition, we found distinct brain regions associated with afferent and efferent cardiac signals. While the “functionally afferent” regions were widely spread over the frontal pole, ACC, PCC, postcentral gyrus, and posterior insula (Figure 3A, blue), the “functionally efferent” regions were focused on the dorsal ACC, anterior insula, and thalamus (Figure 3A, orange). These functionally derived regions were also anatomically identified as parts of the central autonomic network (Benarroch, 1993; Cechetto and Shoemaker, 2009). While the afferent and efferent regions were segregated for younger adults (Figure 3A), these two types of regions overlapped with each other for older adults (Figure 3B). In addition, Granger causality analysis indicated that, for most younger adults, BOLD activity in the afferent regions (Figure 3A blue) predicted BOLD activity in the efferent regions (Figure 3A orange), not the other direction. However, there was no difference between the afferent-to-efferent and efferent-to-afferent predictions in BOLD activity for older adults.

Functionally afferent regions (Figure 3A) appeared along the pathways of relaying incoming cardiac signals to the posterior insula (Nieuwenhuys, 2012). As we predicted, the afferent regions mostly overlapped with the interoceptive regions (Adolfi et al., 2017) including the posterior insula, rostral and dorsal ACC, and somatosensory cortex. Those regions receive cardiovascular signals (Craig, 2003; Khalsa et al., 2009; Oppenheimer and Cechetto, 2016) and relay them to the anterior insula, which is associated with interoceptive awareness and has access to the motor system via dorsal ACC (Craig, 2009). The posterior insula and dorsal ACC showed increased activity when attending to the timing of one’s own heartbeat compared to judging the timing of external sounds (Critchley et al., 2004), suggesting their role in representing visceral sensations. Another study involving both the interoceptive and arousal tasks reported that the mid insula and dorsal ACC were activated in both perception and arousal conditions and speculated that the insula is forwarding the afferent heartbeat information to the ACC to control cardiovascular activity (Pollatos et al., 2007). The broadly activated medial PFC regions, covering the frontal pole and rostral/dorsal ACC (Figure 3A), were consistently reported in neuroimaging studies investigating the functional architecture of autonomic activity, mostly involving cognitive, affective, or motor tasks (Beissner et al., 2013; Ruiz Vargas et al., 2016; Thayer et al., 2012). The functionally afferent regions were also found in resting-state fMRI studies. One study found that BOLD activity in the posterior insula and ACC was negatively correlated with high-frequency HRV, indexing parasympathetic influence, and BOLD activity in the medial PFC was positively correlated with heart rate, whose increase reflects less parasympathetic influence (Valenza et al., 2020). Another resting-state study found that activity in the ventromedial PFC was positively correlated with mean peak-to-peak intervals, reflecting parasympathetic activity (Ziegler et al., 2009).

Functionally efferent regions during rest (Figure 3A) included the dorsal ACC, anterior insula, thalamus, dorsal pons, and cerebellum, most of which overlap with emotional arousal pathways (Adolfi et al., 2017; Min et al., 2022) and mental stress-related regions (Critchley et al., 2003). Among the efferent regions, the anterior insula and dorsal ACC are the main cortical nodes of the salience network which filters constantly incoming signals even at rest and amplifies important information to enhance readiness for goal-directed behavior (Menon, 2015). This coactivation of the anterior insula and ACC in our study provides an interesting contrast between the resting-state and task-engaged brains because the two regions tend to show distinct activation patterns in studies involving cognitive and affective tasks. The anterior insula activates in affective and interoceptive tasks implying afferent signaling (Craig, 2009; Critchley et al., 2004), whereas the dorsal ACC appears during cognitive and motor tasks involving efferent control (Botvinick et al., 2004; Critchley et al., 2003). During rest, the two separate regions might be jointly on standby to orient attention to salient stimuli and modulate cardiac activity to rapidly utilize the motor resources to respond to changing environments (Menon and Uddin, 2010).

The afferent regions’ BOLD activity showed a negative correlation with incoming parasympathetic signals (before-TR RMSSD) but the efferent regions’ BOLD activity showed a positive correlation with outgoing parasympathetic signals (after-TR RMSSD). These relationships do not appear perfectly in line with prior findings. The efferent regions that showed a positive correlation with parasympathetic signals in the current study (Figure 3) were also reported in physical arousal studies that suggested a positive correlation with sympathetic signals (Critchley, 2005), counteracting parasympathetic activity. Rather than a negative correlation in the afferent region, prior studies suggest a positive correlation between afferent signals and activity in the insula and anterior cingulate cortex as the regions were activated during interoceptive tasks involving afferent heartbeat signals (Critchley et al., 2004; Pollatos et al., 2007).

The inconsistencies between the current and prior findings might be due to variations in the tasks. Parasympathetic control regions (i.e., parasympathetic efferent activity) differed across cognitive, affective, and somatosensory-motor tasks (Beissner et al., 2013, Figure 3). Even so, positive correlations in our efferent regions (e.g., the anterior insula, thalamus, and dorsal ACC) were robustly seen in prior studies involving task scans and HRV (Ruiz Vargas et al., 2016, Figure 3). The negative correlations between RMSSD and activity in the afferent regions can be better explained by passive tasks approximating resting state. By using a lower body negative pressure task which did not require volitional effort, one study showed that a decrease in baroreceptor afferent activity (i.e., reduced afferent parasympathetic signaling) was associated with increased activity in the posterior insula but decreased activity in the anterior insula, amygdala, and ACC (Kimmerly et al., 2005). This finding that diminished afferent parasympathetic signaling increased posterior insula activity is consistent with our results. In our study, when before-TR RMSSD (i.e., afferent parasympathetic signaling) was decreased, the afferent regions including the posterior insula increased BOLD activity. This inverse correlation is also seen in a resting-state study that HRV was negatively correlated with BOLD activity in wide cortical areas involving the posterior insula and dorsal ACC (Valenza et al., 2019). In addition, vagus nerve stimulation, increasing parasympathetic activity without mental effort, decreased BOLD activity in the ACC, medial prefrontal cortex, and somatosensory cortex of depressed participants (Nahas et al., 2007).

Taken together, our findings suggest a negative feedback loop where the afferent regions (e.g., the posterior insula) increase BOLD activity in response to decreased parasympathetic activity and activate the efferent regions (e.g., the anterior insula), which in turn enhance parasympathetic activity. One missing link in prior research was whether BOLD activity in the afferent regions leads to activity in the efferent regions. This relationship was supported by the high positive cross-correlations between BOLD activities in the afferent and efferent regions (Figure 5). The correlations were greater when the afferent regions’ BOLD signal was preceding the efferent regions’ signal than the inverse order for younger adults (Figure 5A). This one-directional relationship was further supported by Granger causality analysis showing that BOLD activity in the afferent regions better predicted BOLD activity in the efferent regions than the inverse for younger adults. The asymmetric cross-correlation and Granger causality were not observed for older adults. Additionally, the negative feedback loop, which counteracts momentary changes, does not preclude the possibility that increased parasympathetic activity reduces the afferent and efferent regions’ BOLD activity, leading to a decrease in parasympathetic activity.

Aging appears to affect the functional organizations of afferent and efferent activity. For younger adults (Figure 3A), the afferent and efferent areas were broad but well segregated such that the two regions did not share areas except only a small proportion of each region (6.6% of the afferent and 8.9% of the efferent regions). In contrast, the separation does not seem maintained with aging as the afferent and efferent regions for older adults shared more spatial locations. For older adults, the afferent clusters did not include the prefrontal and cingulate cortices, and a notable portion of the efferent regions (21.3% of the efferent regions) overlapped with the afferent regions (Figure 3B). This might be due to age-related decrease in BOLD activity and specificity in the aging brain during motor and cognitive tasks (Huettel et al., 2001; Kannurpatti et al., 2010). Spontaneous BOLD activity over low-frequency bands at rest is reduced with aging in the PFC, ACC and PCC region (Hu et al., 2014). Resting-state networks become less distinct and more diffuse with aging as within-network connectivity decreased with aging, while between-network connectivity increased (Damoiseaux, 2017; Geerligs et al., 2015).

Returning to the RMSSD-associated regions during emotion regulation, the significant clusters (Figure 2) are likely to be driven more by afferent activity than by efferent activity because the clusters overlap more with the afferent regions (Figure 3A, blue) and, like these regions during rest, showed a negative correlation with time-varying RMSSD. Among the regions showing increased BOLD activity for decreased time-varying RMSSD during the task (Figure 2), the ACC and anterior insula are the main nodes of the salience network (Menon, 2015). This shows an interesting contrast with our previous finding that participants with greater mean (tonic) resting RMSSD better decreased BOLD activity in the default mode network in response to repeated emotional stimuli during emotion regulation (Min et al., 2024, Figure 4). The prior study (Min et al., 2024) suggested that tonic RMSSD at rest might serve as a trait-like marker which indicates individual differences in emotion regulation ability as reflected in the default mode network’s activity. In contrast, time-varying (phasic) RMSSD during a task might represent task-dependent states in cardiac activity as reflected in the salience network’s activity. We also noticed that the clusters during emotion regulation were predominantly focused on the right and medial structures (Figure 2), while both afferent and efferent regions during resting (Figure 3A) were symmetrical (Supplementary Information).

Before we conclude, we acknowledge the study’s limitations. Our findings rely on BOLD signals, indirect measures of neuronal activity, which can be influenced by both excitatory and inhibitory neuronal signals. Our fMRI results cannot unambiguously determine the complex directions of afferent and efferent circuits. Besides the correlational nature of fMRI findings, the spatial (3 mm isotropic) and temporal resolutions (2.4 seconds) are insufficient to capture the millisecond signaling of micrometer-sized neurons. We used PPG signals, detecting heart beat through blood volume changes in finger vessels, to compute RMSSD, a time-domain measure reflecting parasympathetic activity. Electrocardiogram, directly detecting cardiac muscle activity, may provide a more accurate measure of cardiac activity, and high-frequency power may be less noise-susceptible. Smaller clusters in older adults (Figure 3) might be attributed to the smaller sample size and reduced variability in RMSSD values in older adults (Table S1). Our data were collected while lying-down inside a scanner, limiting its generalizability to ambulatory real-world scenarios. Above all, our study assumes that RMSSD before TRs affect brain BOLD activity and RMSSD after TRs are affected by brain BOLD activity. Our findings should be interpreted with caution because a temporally preceding event does not necessarily *cause* a subsequent event. Despite these limitations, our study presents a straightforward approach to address the long-standing inquiry into the in vivo afferent and efferent distinctions of the human brain and suggests a negative feedback loop between the brain and the heart.

## Acknowledgements

We thank our research assistants for their help with data collection: Linette Bagtas, Akanksha Jain, Divya Suri, Sophia Ling, Michelle Wong, Yong Zhang, and Gabriel Shih.

## Availability of data

The data underlying this article are available in https://openneuro.org/datasets/ds003823.

