## Supplementary Figures for "Distinct Brain Systems Support Afferent and Efferent Autonomic Activity"

for

1 University of Southern California, Los Angeles, USA
2. Max Planck Institute for Human Development, Berlin, Germany
3. Virginia Polytechnic Institute and State University, Blacksburg, USA
4. Vanderbilt University, Nashville, USA
5. The State University of New Jersey, New Brunswick, USA
6. University of California Irvine, Irvine, USA

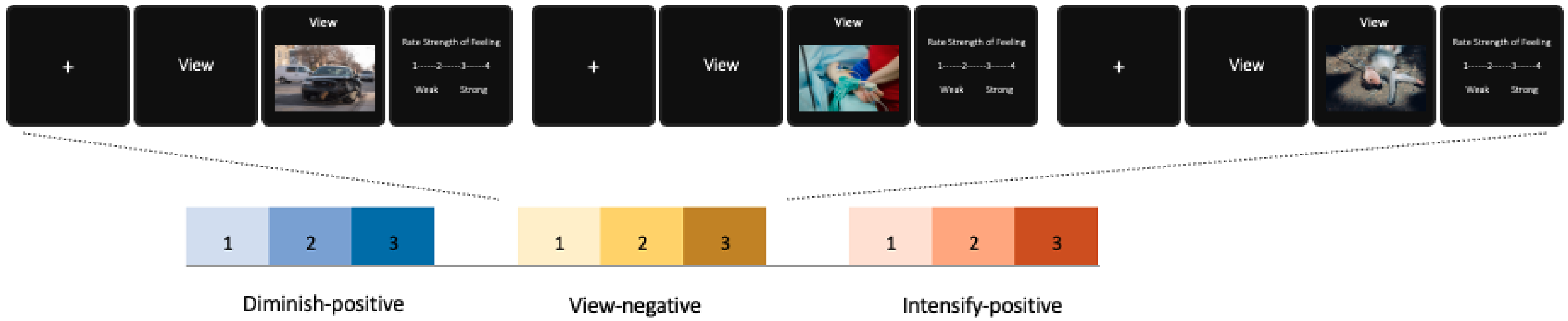


**Figure S1.** Example of the emotion regulation task.The current study employed a mixed block/event-related design (Min et al., 2024). This example shows how three same-condition trials were presented consecutively within a block of the view-negative condition. For the psychophysiological interaction analysis, we used the 36-second block periods of each task condition as a psychological regressor. The event trials of the emotion regulation task were organized in a block-wise manner such that three events of the same condition were contained in each block and separated by a fixation cross lasting for a varying interval. The two intervals separating three events within each block summed to 4 seconds. The 36-second-long blocks were separated by a 5-second fixation cross and presented in a pseudorandom order such that participants did not experience two blocks with the same instruction and image valence in a row. The blocks consisted of seven conditions: view-negative, view-positive, view-neutral, diminish-negative, diminish-positive, intensify-negative, and intensify-positive (six trials per condition). We selected six sets of images with the same average valence ($M_{negative}$ = 2.3, $M_{neutral}$ = 5.0, $M_{positive}$ = 7.2) and same arousal scores ($M_{negative}$ = 5.4, $M_{neutral}$ = 2.8, $M_{positive}$ = 5.4) from the International Affective Picture System (Bradley and Lang, 2017). The standard deviations of image ratings in the six sets were on average ($SD_{negative}$ = 0.3, $SD_{neutral}$ = 0.1, $SD_{positive}$ = 0.3) for valence and ($SD_{negative}$ = 0.3, $SD_{neutral}$ = 0.4, $SD_{positive}$ = 1.0) for arousal. Each set consisted of 18 negative, 18 positive, and 6 neutral pictures, and the images were shuffled within each valence category.

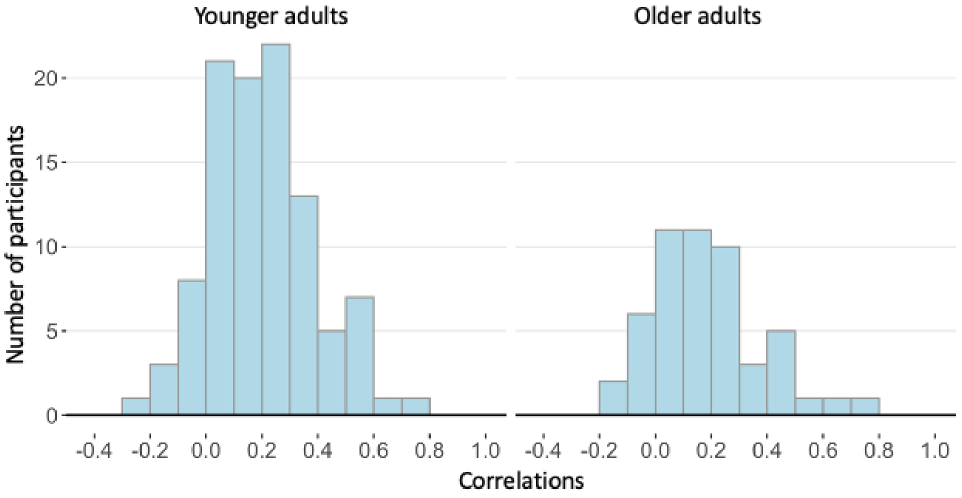


**Figure S2.** Distribution of correlation coefficients between before-TR RMSSD and after-TR RMSSD. To check for high correlations between before-TR and after-TR RMSSD series, which could potentially undermine the validity of the results, we calculated correlations of the before-TR and after-TR RMSSD time series for each participant and ran a one-sample *t*-test on the correlation against 0.5. We also computed the temporal mean and range of the before-TR RMSSD and after-TR RMSSD time series (175 timepoints) within each participant and reported group statistics of the temporal mean and range for younger and older adults (Supplementary Table 1). One-sample t-test against correlation value 0.5 showed that the correlation values were significantly lower than 0.5 for both age groups, $t(101) = -16.36$, $p = 1.88 \times 10^{-30}$ for younger adults and $t(50) = -11.68$, $p = 3.40 \times 10^{-16}$ older adults. The mean and standard deviation of correlation coefficients were 0.199 and 0.186 for younger adults and 0.185 and 0.193 for older adults. The maximum and minimum values were 0.731 and -0.254 for younger adults and 0.729 and -0.185 for older adults.

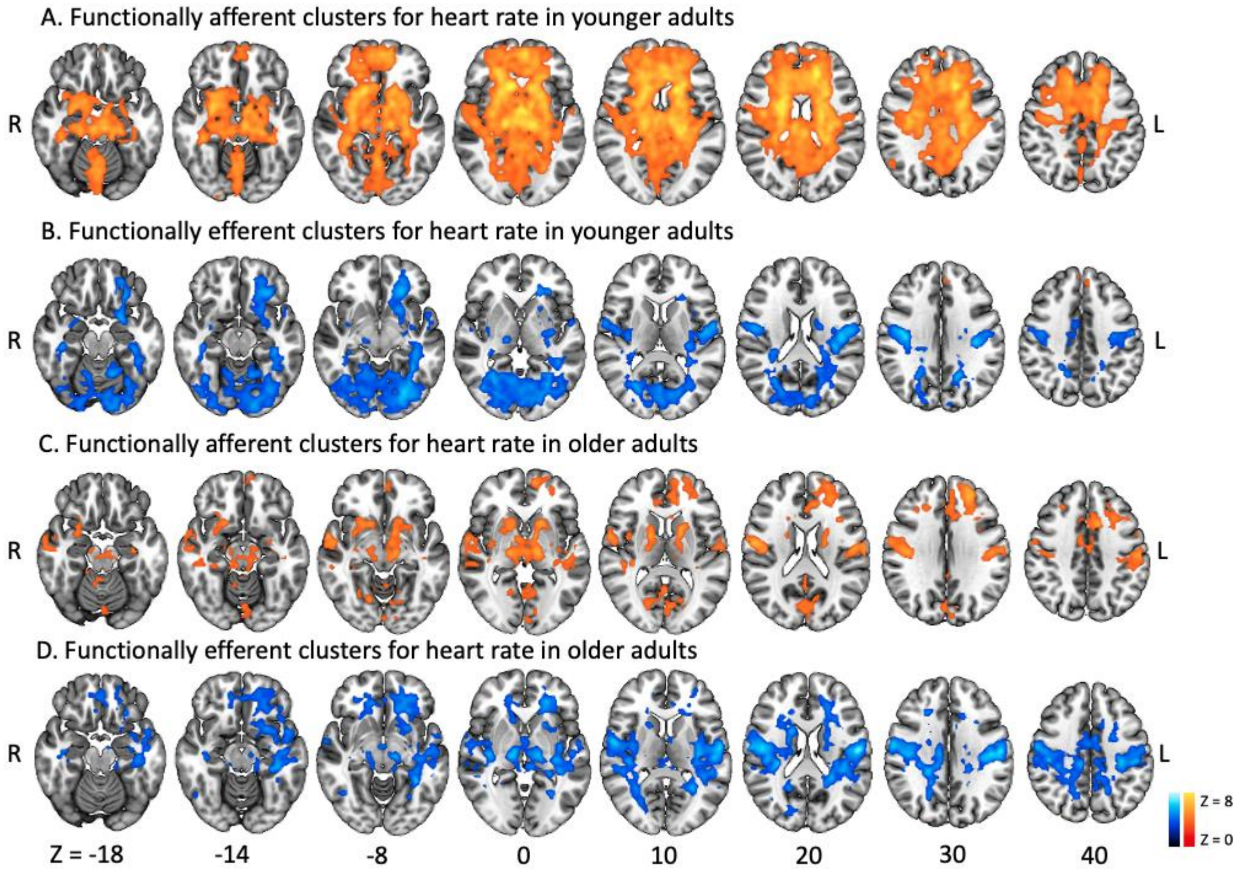


**Figure S3.** Brain regions correlated with heart rate. To confirm that the study results are HRV-specific rather than driven by heart rate itself, we repeated the same whole-brain analyses using before-TR and after-TR heart rates as regressors. Analysis in Figure 3 was repeated with heart rate instead of RMSSD values. Red clusters reflect positive correlations, blue clusters reflect negative correlations. Unlike the neural correlates of RMSSD values (Figure 3), there were no negative but positive correlations with before-TR heart rate in younger adults (A) and older adults (C). There were clusters showing positive (red) and negative (blue) correlations with after-TR heart rate in younger adults (B). There were no positive but negative correlations with after-TR heart rate (D).

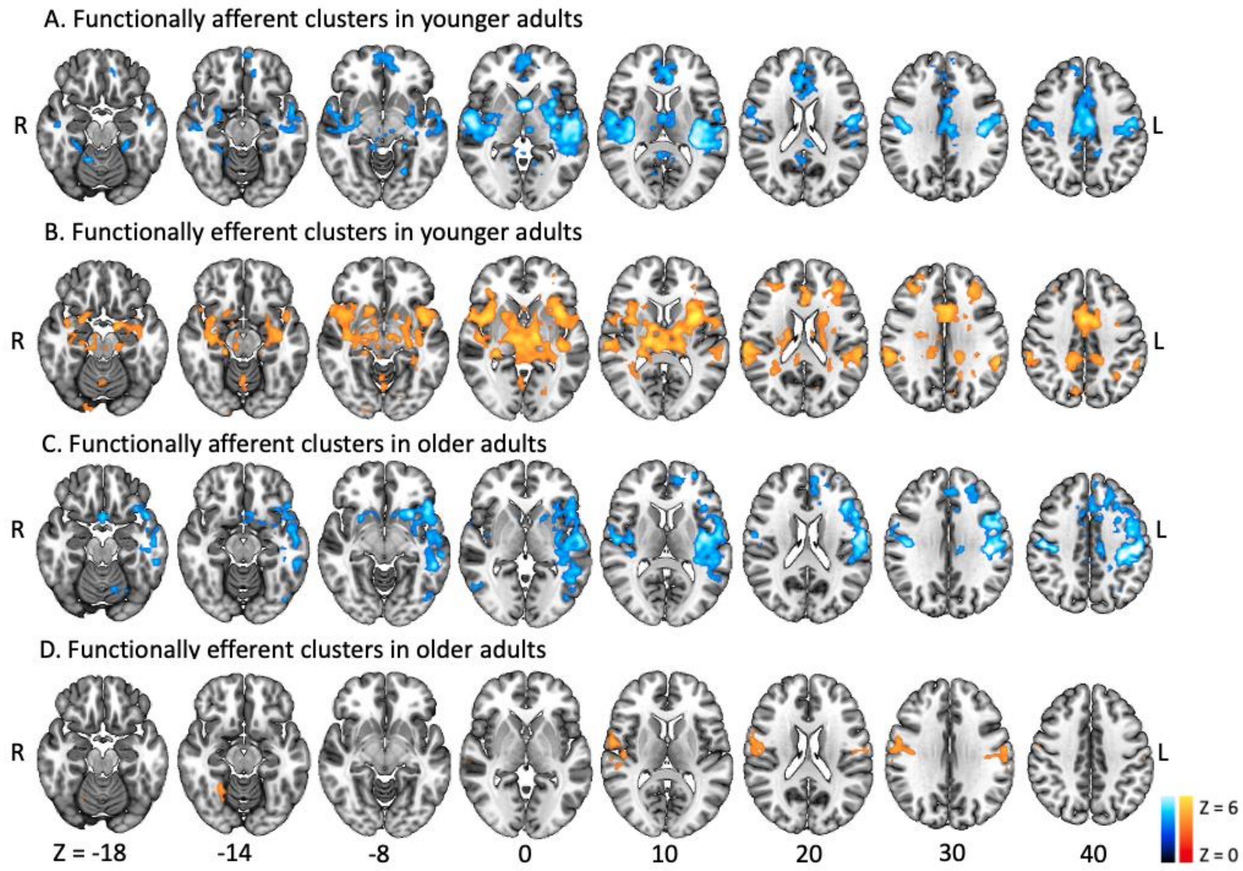


**Figure S4.** Brain regions correlated with either before-TR or after-TR RMSSD. To ensure that the before-TR and after-TR RMSSD time series separately result in similar BOLD activity, we also repeated the whole-brain analyses by having just one regressor in the model. We examined whether similar clusters to Figure 3 would show up when having only one regressor (either before-TR RMSSD or after-TR RMSSD time series) in two simple-regression models rather than having two regressors (both before-TR and after-TR RMSSD time series) in one multiple-regression model (Figure 3). Two separate simple regression models showed similar significant clusters to one multiple regression model.

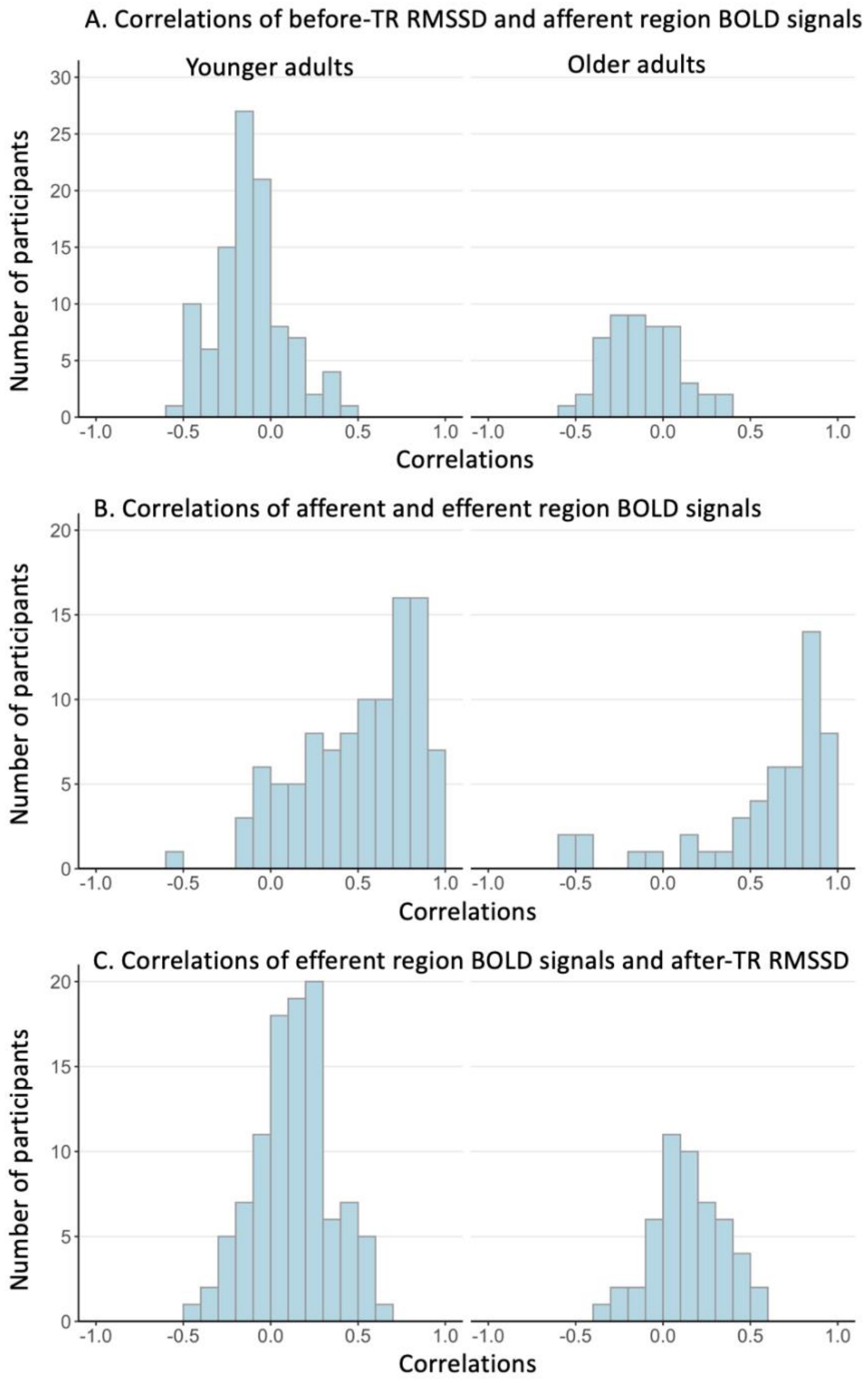


**Figure S5.** Correlations between before-TR RMSSD, afferent regions, efferent regions, and after-TR RMSSD. To further test our proposed feedback loop idea, we additionally ran correlation analyses between BOLD signals from the afferent and efferent clusters (Figure 3) and the two types of RMSSD time series. We examined the relationships across before-TR RMSSD, BOLD signals in the afferent regions (Figure 3), BOLD signals in efferent regions (Figure 3), and after-TR RMSSD. For each individual, we computed correlations between (A) before-TR RMSSD and BOLD signals in afferent regions, (B) between BOLD signals in afferent regions and BOLD signals in efferent regions, (C) between BOLD signals in efferent regions and after-TR RMSSD. Average correlations and standard errors for (A), (B) and (C) were -0.12 (0.02), 0.50 (0.03), 0.13 (0.02) for younger adults and -0.12 (0.03), 0.59 (0.06), 0.15 (0.03) for older adults. To test whether the correlations are aligned with the negative feedback model we proposed, we compared each type of correlation against zero. The negative feedback model suggests negative correlations for (A) and positive correlations for (B) and (C). As predicted by the negative feedback loop, correlation between before-TR RMSSD and BOLD signals in afferent regions (A) was negative, $t(101) = -6.02$, $p = 1.4 \times 10^{-8}$ for younger adults and $t(50) = -4.31$, $p = 3.9 \times 10^{-5}$ for older adults. Correlations between BOLD signals in afferent and efferent regions (B) were positive, $t(101) = 15.30$, $p = 2.4 \times 10^{-28}$ for younger adults and $t(50) = 10.11$, $p = 5.6 \times 10^{-14}$ for older adults. Correlations between BOLD signals in efferent regions and after-TR RMSSD (C) were positive, $t(101) = 6.19$, $p = 6.6 \times 10^{-9}$ for younger adults and $t(101) = 5.37$, $p = 1.0 \times 10^{-6}$ for younger adults.

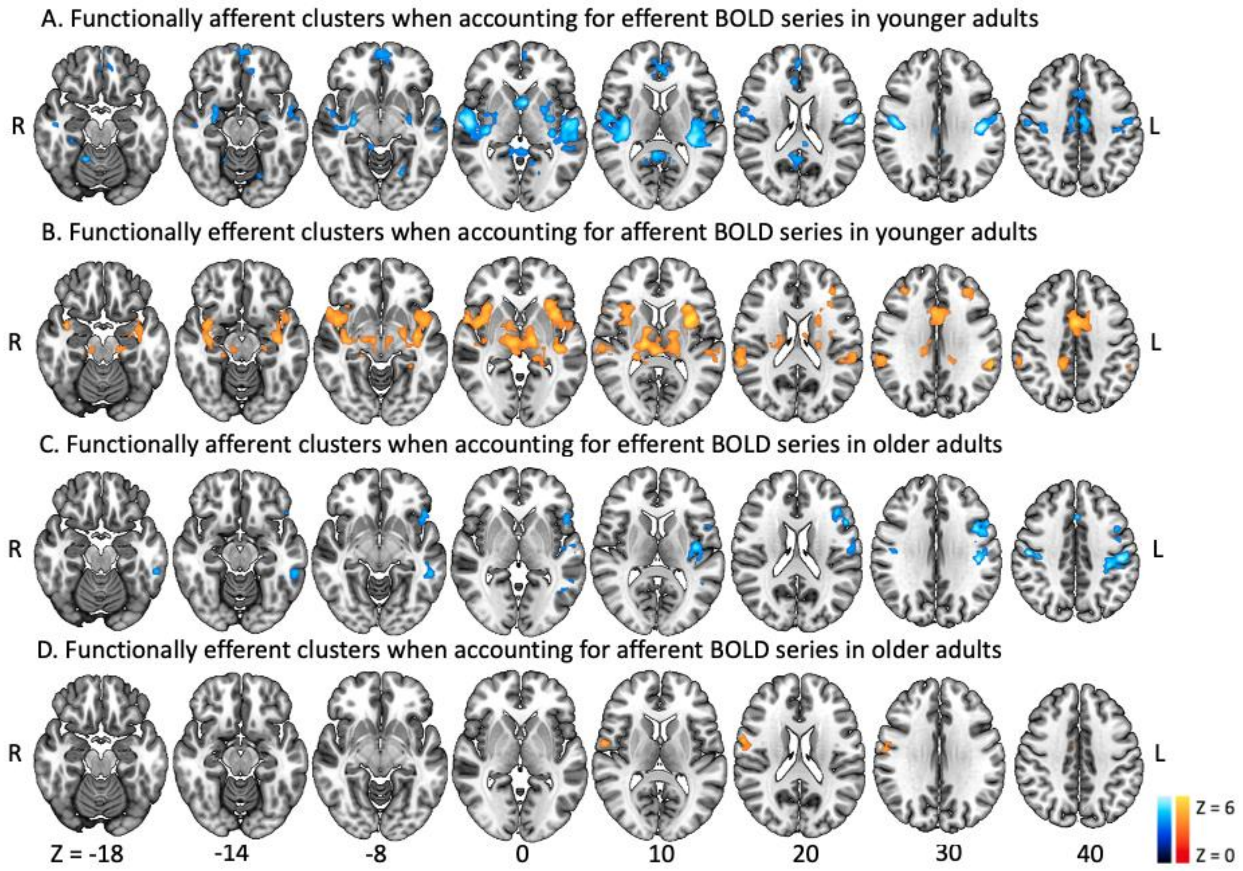


**Figure S6.** Afferent and efferent brain regions when accounting for BOLD signals in the efferent and afferent regions respectively. We examined whether the relationship of before-TR RMSSD and brain BOLD activity in the afferent regions (Figure 3, blue) while accounting for BOLD activity in the efferent regions (Figure 3, orange) remains significant by using before-TR RMSSD time series as a regressor and including BOLD signals from the after-TR RMSSD's neural correlates (Figure 3, orange) as a covariate. The same analysis was repeated with after-TR RMSSD as a regressor and BOLD activity in the afferent regions (Figure 3, blue) as a covariate. To examine whether before-TR brain correlates in Figure 3 remain significant when accounting for after-TR neural correlates, we used before-TR RMSSD time series as a regressor and included BOLD signals from the after-TR RMSSD's brain correlates (extracted from Figure 3 orange regions and averaged across voxels to form one time series) as a covariate. Similarly, to examine the significance of the after-TR brain correlates in Figure 3 when accounting for before-TR brain correlates, we used after-TR RMSSD time series as a regressor and included BOLD signals from the before-TR RMSSD's brain correlates (extracted from Figure 3 blue regions) as a covariate. The significant clusters were similar to the clusters in Figure 3, but smaller in extent.

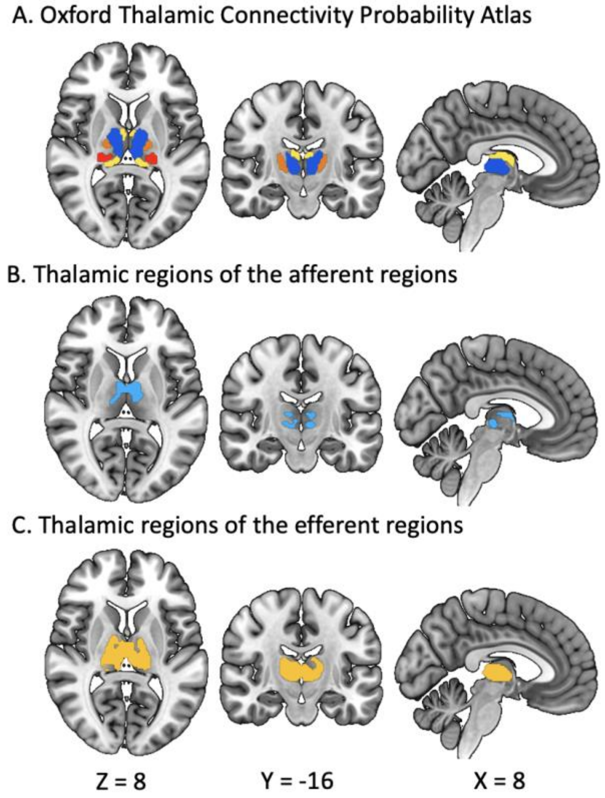


**Figure S7.** Thalamic subregions of the afferent and efferent regions. A shows four subregions with 50% probability based on the Harvard-Oxford Subcortical Structural Atlas. The blue, yellow, orange, and red thalamic subregions are connected to the prefrontal, temporal, premotor, and posterior-parietal cortex respectively for younger adults. B and C show 50%-thresholded thalamic regions of the afferent and efferent regions (Figure 3) respectively for younger adults. The Dice-Sørenson coefficient between the afferent thalamic region (Supplementary Figure 8B) and each thalamic subregion (Supplementary Figure 8A) was 0.38 for prefrontal connection and 0.11 for temporal connection. The Dice-Sørenson coefficient between the efferent thalamic region (Supplementary Figure 8C) and each thalamic subregion (Supplementary Figure 8A) was 0.63, 0.19, 0.17, and 0.16 for prefrontal, temporal, posterior-parietal, and premotor connection respectively.

To examine the nature of thalamic regions associated with before-TR and after-TR RMSSD levels during rest, we used the Oxford Thalamic Connectivity Probability Atlas which divides the thalamus into seven subregions, each of which anatomically connects to occipital, posterior parietal, prefrontal, premotor, primary motor, sensory, or temporal cortices (Behrens et al., 2003). We first segregated the whole thalamus from the afferent and efferent clusters (Figure 3A) by using the 50%-thresholded thalamus mask from the Harvard-Oxford Subcortical Structural Atlas and counted the total number of non-zero voxels of the afferent and efferent thalamic regions. We then counted the overlapping voxels between the segregated afferent and efferent thalamic regions and the binarized masks from the 50%-thresholded Oxford Thalamic Connectivity Probability Atlas. We also calculated the Dice-Sørenson coefficient(Carass et al., 2020) and reported the regions with the coefficients greater than 0.10.

We found that broad regions within the thalamus showed correlations with both types of RMSSD time series for younger adults, but there were no such thalamic correlations for older adults (Figure 3). To explore the potential roles of the thalamus in relaying afferent and efferent signals in younger adults, we used the thalamic connectivity atlas to post-hoc examine which cortical regions the thalamic subregions connect to (Supplementary Figure 8A). In total, 426 voxels out of the identified afferent regions belonged to the thalamus, among which 253 voxels (59.4 %) were connected with the prefrontal cortex, 54 voxels (12.7 %) were connected with the temporal cortex, and 58 voxels (13.6%) were connected with the posterior parietal (27), premotor (21), primary motor (6), and sensory cortex (4) (Supplementary Figure 8B). The thalamus took up a larger part of the efferent regions (Supplementary Figure 8C). In total, 1606 voxels in the efferent regions belonged to the thalamus. Among them, 794 (49.4 %), 199 (12.4 %), 166 (10.3 %), 139 (8.7%) voxels were connected to the prefrontal, temporal, posterior-parietal, and premotor cortex respectively, and. 76 voxels (4.7%) were connected to the primary motor (37), and sensory (28), occipital (11) cortex.

Interestingly, BOLD activity in the thalamus and striatum was correlated with afferent and efferent cardiac activity for younger adults (Figure 3A). While studies suggest the thalamus's involvement in respiratory and cardiovascular functions (Goswami et al., 2011; Kimmerly et al., 2005; Motl et al., 2015; Pattinson et al., 2009; Xu et al., 2013), few studies investigated thalamic contribution to heart-brain interactions. While before-TR RMSSD signals were negatively correlated with BOLD activity in the caudate, putamen and thalamus, after-TR RMSSD signals were positively correlated with BOLD activity in the broad areas of the thalamus (Figure 3A). Our post-hoc analysis revealed that more than half of the afferent and efferent thalamic regions showed functional connections with the prefrontal cortex, highlighting a potential thalamic role in conveying information between the heart and prefrontal regions. Prior studies suggest that the thalamus plays a central role in maintaining the integrity of the central autonomic activity as its volume decrease was associated with reduced baroreceptor sensitivity (De Looze et al., 2020) and reduced cardiorespiratory fitness in multiple sclerosis patients (Motl et al., 2015). Our findings of HRV-associated activity in the caudate and putamen are consistent with prior findings that the gray matter volumes of the caudate and putamen were negatively correlated with high-frequency HRV (Wei et al., 2018), and enhancing dopaminergic activity with methylphenidate administration increased heart rate and blood pressure in humans (Volkow et al., 2003).

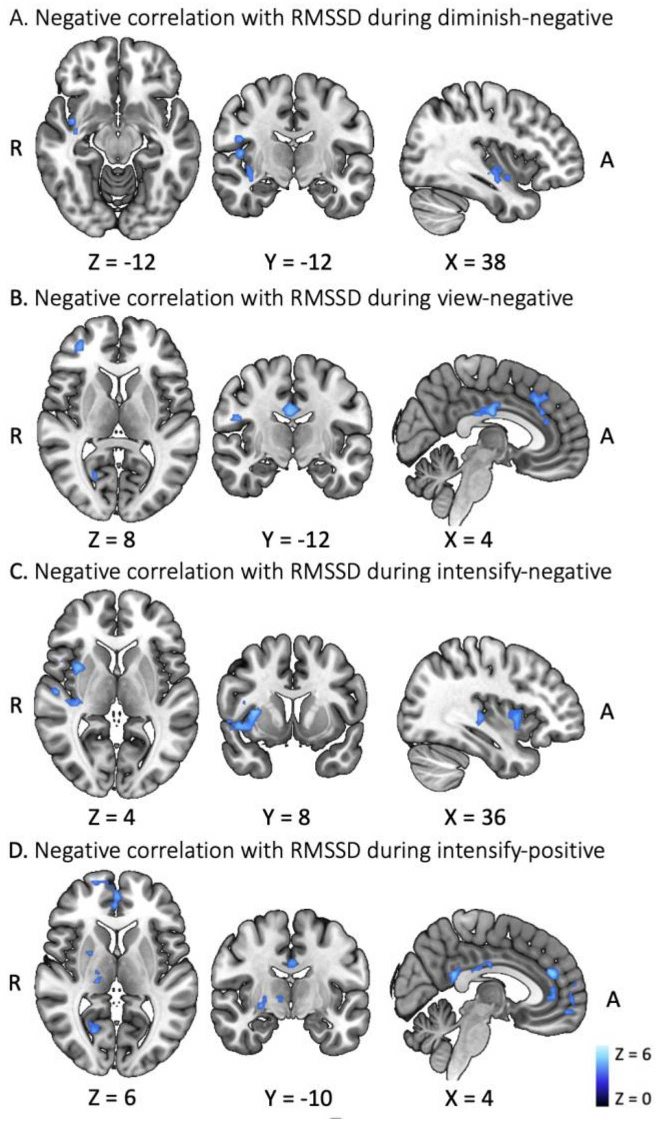

**Figure S8.** Brain regions correlated with RMSSD during regulating negative and positive emotions. For younger adults, BOLD activity in the blue regions was negatively correlated with RMSSD during diminishing negative emotion (A), viewing negative images (B), intensifying negative emotion (C), and intensifying positive emotion (D). There were no positive correlations for younger adults. Neither negative nor positive correlations were found for older adults.

We found BOLD activity in the insula and central opercular cortex was negatively correlated with RMSSD when down-regulating negative emotions (Supplementary Figure 2A). When viewing negative images, activity in the ACC, paracingulate gyrus, middle frontal gyrus and frontal pole showed negative correlations with RMSSD (Supplementary Figure 2B). When up-regulating negative images, RMSSD was negatively correlated with activity in the insula, Heschl's gyrus, and precentral gyrus (Supplementary Figure 2C). There were no significant correlations for down-regulating and viewing positive emotion. When up-regulating positive emotion, activity in the ACC, paracingulate gyrus, frontal pole, PCC, thalamus, putamen, and pallidum were negatively correlated with RMSSD (Supplementary Figure 2D). Of note, all the relationships were predominantly in the right hemisphere during emotion regulation. None of the conditions showed significant clusters for older adults. When we contrasted older adults against younger adults, we found significant clusters including the lingual gyrus and intracalcarine cortex which were driven by the stronger negative correlation of RMSSD in younger than in older adults.

**Discussion S1:** Emotion regulation and brain laterality

We also noticed that the clusters during emotion regulation were predominantly focused on the right and medial structures (Figure 2), while both afferent and efferent regions during resting were not lateralized but rather symmetrical (Figure 3A). The right hemisphere dominance with respect to cardiac activity has been frequently reported in prior studies which investigated the brain substrates of heart rate control (Craig, 2005; Hagemann et al., 2003; Schulz, 2016; Thayer and Lane, 2009). This asymmetry also seems aligned with a group of studies which suggest that the right hemisphere is dominantly associated with affective processing (Borod et al., 1998; Gainotti, 2019; Hartikainen, 2021). One interesting contrast is that our previous study showed left-hemisphere dominance during emotion up-regulation (Min et al., 2022, Figure 3), unlike the right-hemisphere dominance among RMSSD-correlated regions (Figure 2). Because intensifying emotion involves both the cognitive process of amping up emotional intensity as well as the state of arousal resulted from the process, the left-dominant activation during emotion up-regulation might come from a mixed effect of increased attention and arousal as opposed to the parasympathetic influence on the right hemisphere (Figure 2).

**Table S1.** Descriptive statistics of heart rate and RMSSD

| Younger adults (N=102) | Mean | Min | Max | Standard error | 95% CI lower bound | 95% CI upper bound |
|---|---|---|---|---|---|---|
| Within-subject mean of before-TR heart rate (bpm) | 69.285 | 47.273 | 94.546 | 0.984 | 67.333 | 71.237 |
| Within-subject mean of after-TR heart rate (bpm) | 69.252 | 47.287 | 94.334 | 0.985 | 67.299 | 71.206 |
| Within-subject mean of before-TR RMSSD (sec) | 0.050 | 0.012 | 0.197 | 0.003 | 0.044 | 0.055 |
| Within-subject mean of after-TR RMSSD (sec) | 0.049 | 0.012 | 0.198 | 0.003 | 0.044 | 0.055 |
| Within-subject Range of before-TR heart rate (bpm) | 13.389 | 3.661 | 26.401 | 0.491 | 12.414 | 14.364 |
| Within-subject Range of after-TR heart rate (bpm) | 13.506 | 3.462 | 26.350 | 0.489 | 12.535 | 14.476 |
| Within-subject Range of before-TR RMSSD (sec) | 0.074 | 0.014 | 0.234 | 0.005 | 0.064 | 0.083 |
| Within-subject Range of after-TR RMSSD (sec) | 0.074 | 0.014 | 0.238 | 0.005 | 0.065 | 0.084 |
| **Older adults (N=51)** | **Mean** | **Min** | **Max** | **Standard error** | **95% CI lower bound** | **95% CI upper bound** |
| Within-subject mean of before-TR heart rate (bpm) | 67.294 | 48.444 | 100.789 | 1.500 | 64.280 | 70.307 |
| Within-subject mean of after-TR heart rate (bpm) | 67.286 | 48.468 | 100.682 | 1.501 | 64.272 | 70.300 |
| Within-subject mean of before-TR RMSSD (sec) | 0.024 | 0.002 | 0.096 | 0.002 | 0.019 | 0.028 |
| Within-subject mean of after-TR RMSSD (sec) | 0.024 | 0.002 | 0.094 | 0.002 | 0.019 | 0.028 |
| Within-subject Range of before-TR heart rate (bpm) | 8.261 | 2.511 | 21.936 | 0.580 | 7.095 | 9.426 |
| Within-subject Range of after-TR heart rate (bpm) | 8.197 | 2.431 | 22.557 | 0.572 | 7.049 | 9.346 |
| Within-subject Range of before-TR RMSSD (sec) | 0.044 | 0.003 | 0.187 | 0.005 | 0.034 | 0.055 |
| Within-subject Range of after-TR RMSSD (sec) | 0.042 | 0.003 | 0.192 | 0.005 | 0.032 | 0.053 |

*Note*. To examine whether heart rate and RMSSD showed reasonable fluctuations over time within each subject, we computed the temporal mean and range of heart rate and RMSSD over the 175 timepoints for each participant and obtained group statistics of the within-subject means and ranges for younger and older adults. On average, the within-subject mean and range of RMSSD for older adults appear to be lower than those for younger adults, which might have contributed to less significant clusters for older adults (Figure 3B).
